# PARSEbp: Pairwise Agreement-based RNA Scoring with Emphasis on Base Pairings

**DOI:** 10.1101/2025.10.13.682106

**Authors:** Sumit Tarafder, Debswapna Bhattacharya

## Abstract

**Motivation:** High-fidelity scoring of RNA 3D structures remains a major challenge in RNA structure prediction and conformational sampling. While single-model methods for scoring RNA structures can capture individual structural features, they fail to capture the broader structural consensus within a conformational ensemble, limiting their effectiveness in ranking and model selection.

**Results:** We present PARSEbp, a fast and effective multi-model RNA scoring method that integrates pairwise structural agreement across the conformational ensemble with base pairing consistency. By leveraging both alignment-based global structural agreement at the 3D level and base pairing consistency at the 2D level, PARSEbp efficiently constructs a consensus similarity matrix from which per-structure accuracy scores are computed. Tested on 16th Critical Assessment of Structure Prediction (CASP16) RNA targets, PARSEbp significantly outperforms existing single- and multi-model RNA scoring functions, including traditional statistical potentials, state-of-the-art deep learning methods, and consensus-based approaches, as well as a baseline variant of PARSEbp without the emphasis on base pairings, across a wide range of complementary assessment metrics.

**Availability:** PARSEbp is freely available at https://github.com/Bhattacharya-Lab/PARSEbp.

## 1 Introduction

The three-dimensional (3D) structure of RNA plays a central role in determining its catalytic activity, regulatory capacity, and molecular interactions [1, 2]. However, unlike proteins, RNAs are structurally flexible and can adopt diverse conformations stabilized by canonical and non-canonical base pairs, long-range interactions, and interactions with other molecules [3, 4, 5, 6]. Due to the inherent flexibility of RNA molecules, ensemble-based approaches for RNA 3D structure prediction are crucial to capture the range of conformations plausible for a given sequence [7]. Recent advances in computational modeling of RNA 3D structures involving both deep learning-based methods [8, 9, 10, 11] and stochastic energy- or template-based approaches [12, 13] enable the generation of a large pool of candidate 3D structures for a given sequence. Despite the progress, automated prediction of RNA 3D structures still lags behind the human-expert modeling groups in the recent community-wide blind assessments [14, 15, 16], largely due to data scarcity, structural complexity, and weak evolutionary signal of RNA sequences [17]. To render the computational ensembles practically useful, RNA scoring methods are essential to identify high-quality models from misfolded alternatives present in the ensemble to improve the accuracy of RNA 3D structure prediction [18, 19].

Existing approaches for 3D RNA scoring can be broadly divided into two categories: single-model and multi-model methods. Single-model scoring methods evaluate each candidate structure independently without considering the alternative conformations available in the ensemble. These methods rely on either knowledge-based statistical potentials [20, 21, 22, 23] or deep learning architectures [24, 25, 26, 27] to estimate specific structural quality scores. However, all single-model scoring methods rely on experimental structures, either as supervised signal to train their architecture or fine-tune the statistical potential parameters, thus limited to the paucity of the available RNA 3D structures in Protein Data Bank (PDB) [28]. Additionally, such methods often fail to capture the broader consensus within a structure ensemble, limiting their effectiveness in ranking and model selection. In contrast, multi-model scoring methods can exploit the agreement across the ensemble of candidate structures to infer individual model quality in an unsupervised manner, without the need for experimental structures to train or fine-tune the parameters. A recent method RNAtive [29], the only existing publicly available multi-model RNA scoring method in the literature, derives consensus base pairs by identifying recurrent patterns across RNA 3D folds to assign individual quality scores. However, RNAtive does not perform an explicit all-vs-all pairwise comparison across the ensemble, limiting the ability to capture the full spectrum of structural agreement. This motivates the need to develop an accurate and computationally efficient multi-model RNA scoring method that can integrate both pairwise global structural similarity and base pairing fidelity across the entire ensemble to deliver robust quality estimates.

In this work, we present PARSEbp, a pairwise multi-model RNA scoring method that integrates global 3D structural similarity with 2D base pairing consistency. PARSEbp computes pairwise TM-scores [30] across the structural ensemble to capture global fold similarity with additional emphasis from pairwise Interaction Network Fidelity (INF) scores [31], which quantifies the agreement of canonical and non-canonical base pairs between structures. These measures are combined into a consensus similarity matrix, from which per-structure quality scores are computed by averaging the agreement with all other models in the structural ensemble. Benchmarking on 25 RNA targets from the recently concluded CASP16 challenge, PARSEbp achieves the highest global Spearman’s correlation (0.92) while attaining the lowest top-1 loss of 0.05, demonstrating its effectiveness in top model selection from the diverse structural ensemble in CASP16. By unifying pairwise agreement at the 3D and 2D levels, PARSEbp provides a fast, accurate, and consensus-driven quality estimation tool, requiring only a few minutes to score an entire structural ensemble for a typical RNA of 100 nucleotides.

## 2 Materials & Methods

Given a set of *N* RNA 3D structures *S* = *{S*_1_, *S*_2_, …, *S*_*N*_ *}* as input, we evaluate the structural quality of the decoys by combining pairwise 3D structural similarity with base pair (2D) consistency among the structures in the pool. Specifically, for each ordered pair of structures (*S*_*i*_, *S*_*j*_ ) with *i* ≠ *j*, we compute the pairwise similarity score *T*_*ij*_ as follows:

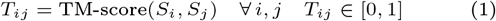

This results in a complete *N × N* pairwise similarity matrix *T* = [*T*_*ij*_ ] which encodes the global structural similarity in terms of TM-score [30] across the ensemble. To guide this global pairwise similarity, we further assess the base pairing consistency among the structures by extracting 2D base pair map 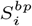 from each 3D structure using MC-annotate [32], a widely used method for RNA base pair annotation. For every pair of structures (*S*_*i*_, *S*_*j*_ ), we compute the pairwise Interaction Network Fidelity (INF) score [31], which quantitatively measures the overlap of base pairing patterns as a degree of agreement between the two structures. INF is defined as the geometric mean between precision and recall and is calculated in the following way:

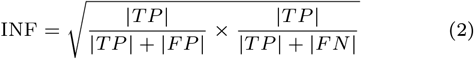

Here, true positives (*TP* ) denote overlapping interaction pairs between structures (*S*_*i*_) and (*S*_*j*_ ) while false positives (*FP* ) and false negatives (*FN* ) denote the set of interactions only found in either (*S*_*i*_) or (*S*_*j*_ ), respectively. The pairwise base pair consistency *I*_*ij*_ between two maps 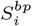 and 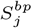, yielding the pairwise base pair similarity matrix *I* = [*I*_*ij*_ ] is defined as:

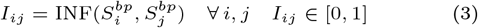

We combine the pairwise 3D and 2D agreements by taking an element-wise multiplication between the two similarity matrices *T* and *I* as follows:

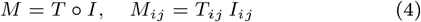

This yields the consensus matrix *M* = [*M*_*ij*_ ], which captures both the global 3D similarity between structures and the extent to which this similarity is reinforced by consistent base pairing interactions. The quality score *Q*_*i*_ for each structure *S*_*i*_ is defined as the mean agreement with all other structures in the ensemble:

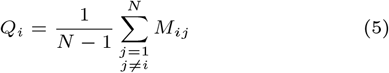

The resulting set of scores *Q* = *{Q*_1_, *Q*_2_, …, *Q*_*N*_ *}* provides a quantitative self-assessment of the structures, where higher scores correspond to structures consistently supported by both 3D structural similarity and 2D base pairing patterns across the ensemble. To ensure scalability for long RNA sequences and large conformational ensembles, all pairwise computations are parallelized and executed in batches.

We evaluate our method PARSEbp against a broad set of both single-model and multi-model RNA scoring methods. Among the single-model methods, we compare against traditional knowledge-based statistical potentials such as rsRNASP [20], rsRNASP1 [21], cgRNAsp [22], DFIRE-RNA [23] and state-of-the-art deep learning-based approaches, including RNA3DCNN [24], ARES [27], lociPARSE [25] and RNArank [26]. For multi-model methods, we compare PARSEbp against the only available web-based tool RNAtive [29] along with a baseline variant of PARSEbp without the emphasis on base pairings. Our benchmark set contains 4,750 decoy 3D structures across 25 RNA targets with sequence length ≤ 500, as listed in **Supplementary Table S1**. Targets with length *>* 500 are omitted from evaluation, as CASP16 results indicate that they predominantly yielded low-quality decoys in the ensemble [16]. Ground truth scores of all the structures are obtained from the official CASP16 assessment, following the CASP16 composite evaluation formula [16] proposed by the assessors, defined as the weighted combination of three structural similarity measures: TM-score [30], Local Distance Difference Test (lDDT) [33] with stereochemical checks enabled, and GDT_TS [34], as follows:

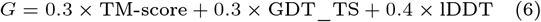

The scoring performance of various methods is evaluated using global Pearson’s correlation (*r*), Spearman’s rank correlation (*ρ*), and root mean squared error (RMSE) between predicted and ground truth scores, along with per-target average *r, ρ*, and loss (top-1 loss). which is defined as the absolute difference between the ground truth of the top-ranked structure and that of the best structure for each target.

## 3 Results

**Table 1** summarizes the performance of PARSEbp against competing single-model and multi-model RNA scoring methods on 25 CASP16 RNA targets using the composite ground truth metric *G* (Eq. 6). At the global level, PARSEbp achieves the highest Pearson’s correlation (*r* = 0.91) and Spearman’s correlation (*ρ* = 0.92) with respect to ground truth scores while yielding the lowest root mean square error (Error = 0.11). On a per-target basis, our method consistently delivers superior correlations and the lowest average loss compared to all other methods. PARSEbp exhibits a remarkable performance advantage compared to single-model approaches. Knowledge-based statistical potentials such as rsRNASP, rsRNASP1, cgRNAsp, and DFIRE-RNA achieve only modest performance, with global correlations not exceeding *r* = 0.22 and substantially higher errors, underscoring the challenge of generalizing from limited reference states. Deep learning–based methods perform comparatively better, with RNArank attaining the highest global Pearson’s correlation among single-model approaches (*r* = 0.72) and lociPARSE achieving the highest per-target average Pearson’s correlation (*r* = 0.70). In terms of per-target average loss, lociPARSE and RNA3DCNN both reach 0.09, representing the best performance within the single-model methods. Nonetheless, PARSEbp surpasses all single-model approaches in both global and per-target evaluations by large margins, achieving the highest per-target average Pearson’s correlation (*r* = 0.85) and the lowest top-1 loss (0.05). These results demonstrate the reliability of PARSEbp in consistently delivering top-notch performance across various complementary assessment metrics. Detailed per-target results are available in **Supplementary Table S2**.

**Table 1.**
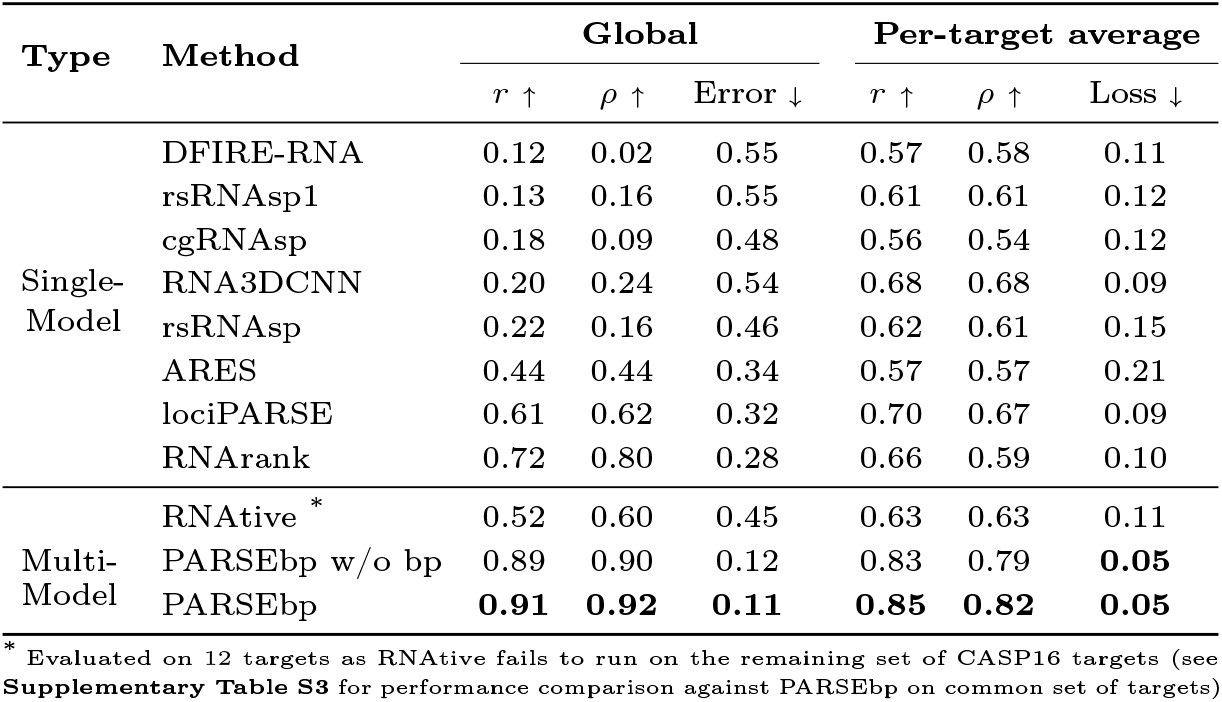
Performance on 25 CASP16 RNA targets (4,750 decoys) based on the CASP16 official assessment metric as the ground truth, sorted in increasing order of global Pearson’s *r*. Values in bold indicate the best performance.

When considering multi-model RNA scoring approaches, PARSEbp is compared against RNAtive, the only other multi-model method currently available in the literature. However, RNAtive is not only limited by ensemble size but also fails on roughly half of the CASP16 targets, particularly for targets dominated by low-quality structures, necessitating additional filtering based on clash score, bond lengths, or angle geometry to proceed, and ultimately making it unsuitable for evaluation across the full set of targets. On the reduced set of 12 targets where RNAtive runs successfully, its performance remains modest with a global error of 0.45 and a per-target average loss of 0.11, lower than even the state-of-the-art single-model approaches. To ensure fairness, we benchmark PARSEbp on this reduced set and report the results in **Supplementary Table S3**. On this reduced benchmark of common set of targets, PARSEbp consistently outperforms RNAtive, achieving a global Pearson’s correlation of *r* = 0.87 and error of 0.13, much better than RNAtive’s *r* = 0.52 and error of 0.45. At the per-target level, PARSEbp attains a loss of 0.03, far lower than that of RNAtive (0.11). To further assess the contribution of the emphasis on the base pair, we compare PARSEbp with a baseline variant that relies only on pairwise TM-score similarity, without the emphasis on base pairings. As shown in **Table 1**, removing base pairs leads to a noticeable drop in performance in terms of both global and per-target average correlations. The drops in correlations are statistically significant at 95% confidence level as revealed by the Wilcoxon signed-rank test with *P*-value = 0.02 for Pearson’s *r* and *P*-value = 0.03 for Spearman’s *ρ*, confirming that the incorporation of base pair consistency provides a meaningful gain in performance for multi-model RNA scoring. In summary, our method significantly outperforms the existing multi-model RNA scoring approach RNAtive and base pair information meaningfully contributes to the improved performance of PARSEbp.

In terms of computational efficiency, PARSEbp completes ensemble-wide scoring for typical RNA targets of length approximately 100 nucleotides within half a minute. By comparison, RNAtive requires substantially longer runtime (approximately half an hour) even for shorter targets (*<*= 100), highlighting the suitability of PARSEbp for large-scale applications. Runtime analysis on the CASP16 benchmark set, provided in **Supplementary Figure S1**, further confirms that PARSEbp exhibits scalability with increasing sequence length and structural ensemble size without sacrificing accuracy. Collectively, these results demonstrate that PARSEbp is a robust and accurate method for multi-model RNA scoring.

## 4 Conclusion

We present PARSEbp, a fast, accurate, and reliable multi-model RNA scoring method that unifies pairwise global structural similarity at the 3D level and base pairing consistency at the 2D level. By explicitly performing all-vs-all alignment-based comparisons given a structural ensemble with emphasis on base pairings, PARSEbp provides consensus-driven high-fidelity scoring of RNA 3D structures, significantly outperforming all existing single- and multi-model methods in CASP16 test set. PARSEbp should be broadly useful for scoring RNA 3D structures at scale.

## 5 Data Availability

The open-source PARSEbp method is freely available at https://github.com/Bhattacharya-Lab/PARSEbp.

The RNA 3D decoy structures and corresponding ground truth scores for the CASP16 RNA targets have been downloaded from https://predictioncenter.org/download_area/CASP16/predictions/RNA/ and https://predictioncenter.org/casp16/results.cgi?view=targets&tr_type=rna, respectively.

## 5 Acknowledgements

This work was partially supported by the National Institute of General Medical Sciences (R35GM138146 to D.B.) and the National Science Foundation (DBI2208679 to D.B.).

