## Supplementary Information for "PARSEbp: Pairwise Agreement-based RNA Scoring with Emphasis on Base Pairings"

### Supplementary Tables

#### S1 List of targets in CASP16 benchmark set

Table S1: Summary of 25 CASP16 RNA targets of length <= 500 nucleotides and their corresponding number of submitted decoy 3D structures from different participating groups in CASP16 competition (Total 4750 decoy structures).

| ID | Length | Number of decoys |
|---|---|---|
| R1288 | 58 | 193 |
| R1205 | 59 | 187 |
| R1263 | 64 | 217 |
| R1264 | 64 | 206 |
| R1209 | 72 | 213 |
| R1296 | 72 | 230 |
| R1271 | 77 | 173 |
| R1293 | 82 | 214 |
| R1221s3 | 86 | 185 |
| R1224s3 | 86 | 199 |
| R1261 | 89 | 227 |
| R1262 | 89 | 216 |
| R1211 | 90 | 185 |
| R1255 | 124 | 190 |
| R1256 | 127 | 188 |
| R1203 | 134 | 183 |
| R1242 | 205 | 191 |
| R1212 | 247 | 202 |
| R1289 | 284 | 157 |
| R1224s2 | 395 | 196 |
| R1221s2 | 398 | 189 |
| R1248 | 407 | 172 |
| R1254 | 413 | 83 |
| R1241 | 480 | 180 |
| R1291 | 480 | 174 |

#### S2 Detailed per-target average results of PARSEbp in CASP16

Table S2: Per-target average performance of PARSEbp on 25 CASP16 RNA targets in terms of Pearson's correlation ($r$), Spearman's correlation ($\rho$), and top-1 loss. Each row represents the average metrics for every target across its own structural decoy ensemble.

| **ID** | **Pearson's $r$** | **Spearman's $\rho$** | **Top-1 Loss** |
|---|---|---|---|
| R1288 | 0.58 | 0.66 | 0.05 |
| R1205 | 0.74 | 0.72 | 0.02 |
| R1263 | 0.93 | 0.95 | 0.00 |
| R1264 | 0.93 | 0.95 | 0.04 |
| R1209 | 0.67 | 0.77 | 0.00 |
| R1296 | 0.79 | 0.83 | 0.15 |
| R1271 | 0.93 | 0.83 | 0.01 |
| R1293 | 0.68 | 0.71 | 0.21 |
| R1221s3 | 0.91 | 0.79 | 0.06 |
| R1224s3 | 0.92 | 0.85 | 0.05 |
| R1261 | 0.95 | 0.88 | 0.05 |
| R1262 | 0.95 | 0.88 | 0.05 |
| R1211 | 0.82 | 0.77 | 0.19 |
| R1255 | 0.85 | 0.87 | 0.01 |
| R1256 | 0.76 | 0.77 | 0.05 |
| R1203 | 0.93 | 0.93 | 0.02 |
| R1242 | 0.93 | 0.90 | 0.03 |
| R1212 | 0.80 | 0.83 | 0.08 |
| R1289 | 0.97 | 0.93 | 0.01 |
| R1224s2 | 0.98 | 0.95 | 0.02 |
| R1221s2 | 0.96 | 0.92 | 0.04 |
| R1248 | 0.86 | 0.86 | 0.05 |
| R1254 | 0.43 | 0.25 | 0.03 |
| R1241 | 0.98 | 0.88 | 0.05 |
| R1291 | 0.98 | 0.83 | 0.07 |
| **Average** | **0.85** | **0.82** | **0.05** |

#### S3 Comparison with RNAtive on common set of targets

Table S3: Performance comparison with RNAtive on a common set of 12 CASP16 RNA targets (R1288, R1205, R1203, R1209, R1221s3, R1224s3, R1254, R1255, R1256, R1263, R1264, R1271) based on the CASP16 assessment metric as the ground truth. Values in bold indicate the best performance.

| Method | Global | | | Per-target average | | |
|---|---|---|---|---|---|---|
| | $r \uparrow$ | $\rho \uparrow$ | Error ↓ | $r \uparrow$ | $\rho \uparrow$ | Loss ↓ |
| RNAtive | 0.52 | 0.60 | 0.45 | 0.63 | 0.63 | 0.11 |
| PARSEbp | **0.87** | **0.89** | **0.13** | **0.80** | **0.78** | **0.03** |

### Supplementary Figures

#### S1 Runtime requirement of PARSEbp

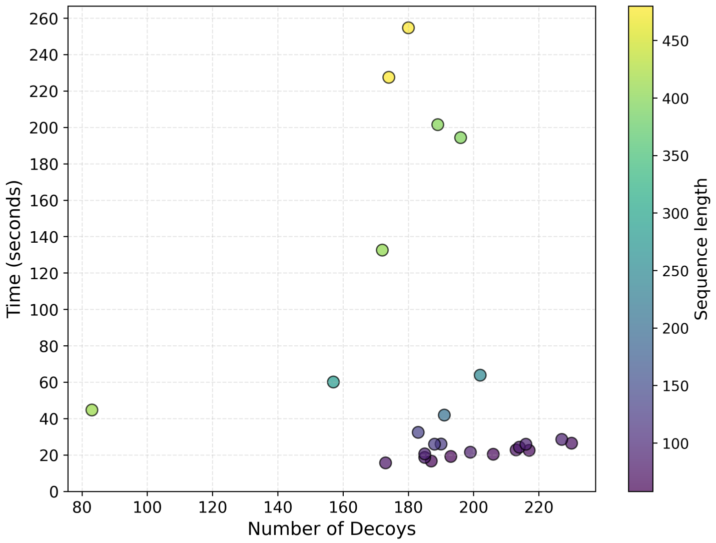

Figure S1: Scatter plot of number of decoys versus runtime (seconds) required for 25 CASP16 targets where the color ramp indicates the increase in sequence length. For typical RNA targets with sequence length of ∼ 100 nucleotides, the runtime is very low (approximately 25-35 seconds) with 50 parallel threads to score the entire ensemble of 3D structures, demonstrating fast and efficient performance, while longer sequences with larger decoy sets naturally require more time.
